# Longitudinal assessment of brain structure and behavior in youth with rapid weight gain: Potential contributing causes and consequences

**DOI:** 10.1101/2022.06.04.494820

**Authors:** Shana Adise, Andrew T. Marshall, Sage Hahn, Shaomin Zhao, Eric Kan, Kyung E. Rhee, Megan M. Herting, Elizabeth R. Sowell

**Affiliations:** Children’s Hospital of Los Angeles, Division on Research on Children, Youth and Families; University of Vermont, Department of Psychiatry; University of California, San Diego, Department of Pediatrics; University of Southern California, Keck School of Medicine, Departments of Population and Public Health Sciences and Pediatrics

**Keywords:** MRI, pediatric obesity, biomarker, eating disorders

## Abstract

**Objective:** Independent of weight status, rapid weight gain has been associated with underlying brain structure variation in regions associated with food intake and impulsivity among pre-adolescents. Yet, we lack clarity on how developmental maturation coincides with rapid weight gain and weight stability.

**Methods:** We identified brain predictors of two-year rapid weight gain and its longitudinal effects on brain structure and impulsivity in the Adolescent Brain Cognitive Development^SM^ Study®. Youth were categorized as Healthy Weight/Weight Stable (WS_HW_, *n*=225) or Weight Gainers (WG, *n*=221, >38lbs); 63% of the WG group were healthy weight at 9-to-10-years-old.

**Results:** A five-fold cross-validated logistic elastic-net regression revealed that rapid weight gain was associated with structural variation amongst 39 brain features at 9-to-10-years-old in regions involved with executive functioning, appetitive control, and reward sensitivity. Two years later, WG youth showed differences in change over time in several of these regions and performed worse on measures of impulsivity.

**Conclusions:** These findings suggest that brain structure in pre-adolescence may predispose some to rapid weight gain and that weight gain itself may alter maturational brain change in regions important for food intake and impulsivity. Behavioral interventions that target inhibitory control may improve trajectories of brain maturation and facilitate healthier behaviors.

## Introduction

Rapid weight gain is defined as abnormal growth within a short period and contributes to childhood obesity risk and exacerbated metabolic consequences.^1–4^ However, apart from infancy, rapid weight gain later in development is poorly understood. The neural mechanisms that may contribute to weight stability versus cause others to experience rapid (or excessive) weight gain independent of their weight status (i.e., healthy weight vs. overweight/obese) remain elusive.

Given the current obesity epidemic in youth,^5^ it is imperative to understand the mechanisms driving weight stability versus rapid weight gain during childhood. The 10-year longitudinal Adolescent Brain Cognitive Development^SM^ Study (ABCD Study®) permits a closer investigation into whether neural differences precede or result from excessive weight gain among 9-to-10-year-old children. Furthermore, we can learn how neural differences correlate with behavior and potentially identify targets of future interventions.

The predictors and long-term effects associated with abnormal rapid weight gain during infancy are well-documented. Correlates of rapid weight gain during infancy include, being born small^6^ or large^7^ for gestational age, prenatal stressors such as substance exposure, preterm delivery, maternal obesity, and undernutrition.^6, 8, 9^ Rapid weight gain during infancy been associated with later obesity risk^10^ and medical comorbidities, such as reduced lung function,^1^ early menarche,^2^ and cardiometabolic risk.^3, 4^ Rapid weight gain during puberty,^11, 12^ while less studied than in infancy, may be more consequential for cardiometabolic health,^13^ highlighting the need for further study in this age range.

Generally, weight gain occurs in response to a surplus of calories (i.e., overeating). Although reasons for overeating and obesity are multifactorial, the brain plays a key role as it controls food intake via homeostatic and hedonic control pathways.^14, 15^ Within this realm, obesity in children has been correlated with altered brain structure,^16^ resting-state functional connectivity,^17, 18^ brain activity during a working memory task,^19^ and altered brain responses in reward and inhibitory control regions to pictures of food,^20, 21^ suggesting that aberrations in brain structure and function may be one reason for overeating. We have previously shown that there are brain regions in 9-to-10-year-old children that predict one-year rapid weight gain,^19^ but it is not known whether weight gain itself may affect trajectories of change in brain structure *because* of rapid weight gain.

The current study assessed the longitudinal relationship between brain structure and weight gain, independent of weight status, in a cohort of youth classified as either Healthy Weight/Weight Stable (WS_HW_) or Weight Gainers (WG) over a two-year period in development. We first assessed if brain regions predictive of one-year weight gain^19^ showed continued structural variation after weight gain onset. However, because rapid weight gain can be temporally sensitive (e.g., some youth may gain weight within one year and stop, while other youth will continue to show continued multiyear rapid weight gain trajectories), we also assessed if there was structural variation at baseline that was predictive of youth who would have sustained, two-year weight gain and how these regions changed after two-years of weight gain. To contextualize extreme weight gain in terms of observable behavior, we investigated neurocognitive metrics focusing on reward and inhibitory control, as deficits in these decision-making processes have been linked to both overeating and obesity.^20–23^ A greater understanding of the relationship between rapid weight gain and brain structure may permit more accurate identification of children at risk for obesity, thereby allowing for interventions to prevent risky eating behaviors before they start.

## Methods

### Study design

Data were curated from the ABCD Study^®^ (3.0 data release), which is a 21-site 10-year longitudinal cohort study aimed to assess neurocognitive development from 9-to-20-years-old. A general overview of the ABCD Study^®^ has been published elsewhere.^24–27^ Here, we focused on anthropometric data from the baseline appointment (9-to-10-years-old) and the one- and two-year follow-up assessments, as well as the neuroimaging data collected at baseline and the two-year follow-up (ages 11-to-12-years-old). Data were available for the entire sample at baseline and one-year follow-up (*n*_baseline_=11,878; *n*_year1_=11,235) and contained half of the participants’ data for the two-year follow-up (*n*_year2_=6,571 youth). Data collection for the baseline assessment occurred between 2016-2018.

### Exclusion criteria

Details of exclusion criteria during screening for participation have been previously published^28^ and include magnetic resonance imaging (MRI) contraindications (e.g., metal implants), not being fluent in English, a history of major neurological disorders (e.g., low functioning autism), premature birth <28 weeks, infant hospitalization >30 days after birth, and disinterest in committing to a longitudinal study.

The current manuscript excluded youth from the analyses if they met the following at any of the time points (e.g., baseline, year 1, or year 2): (1) underweight (according to the Center for Disease Control’s [CDC’s] age-sex-height-weight-specific growth curves^29^); (2) took medications known to alter food intake (e.g., antipsychotics, antidepressants, insulin); (3) met criteria for neurological, psychiatric, or learning disabilities (e.g., attention deficit hyperactive disorder); (4) met diagnostic criteria for eating disorders (e.g., anorexia, bulimia, binge eating disorder) as assessed by the caregiver-reported Kiddie Schedule for Affective Disorders and Schizophrenia (K-SADS)^30^; (5) mislabeled sex-assigned at birth combined with a mismatch with sex-specific pubertal questionnaires or transgendered youth (i.e., due to sex-specific effects on brain function); (6) missing covariate data; (7) height measurement error (e.g., decrease in height over time) (see **Supplemental Materials Table S1** for details); or 8) youth with weight loss to avoid those with restrictive eating habits (e.g., dieting). MRI quality control was performed by the ABCD Study’s^®^ Data Analytics, Informatics, & Resource Center. Tabulated exclusion criteria were provided for the user to apply.

### Anthropometrics

Annually, height and weight were measured twice (and then automatically averaged), by a trained researcher, to the nearest 0.1in and 0.1lb; a third measurement was collected if there was a large discrepancy. Height and weight were converted into BMI (kg/m^2^) and BMI *z*-scores (BMI*z*) and percentiles according to the CDC’s sex-age-heigh-weight-specific cutoffs^29^ per CDC-provided SAS code.

### Weight stability assessment

To evaluate excessive weight gain beyond normative development, youth were split into two groups based on three years of anthropometric data: WS_HW_ and WG. Groups were defined based on clinical cutoffs and weight gain criteria. Clinical cutoffs for weight stability have used BMI *z*-score standard deviation (SD) criterion,^31–34^ in which an SD<0.2 is considered weight stable, and an SD≥0.2 is considered not weight stable (**Fig1A**). Yet, BMI z-scores are poor indicators of weight gain over time^35^ and have several methodological limitations (see Hendrickson et al., (2021)^36^ and Palmer et al., (2021)^37^). Further, the clinical criterion did not capture rapid weight gain (**Fig1B**), so we added a broader weight criterion to better identify youth who were truly WS_HW_ or WG. The WS_HW_ youth had a BMI *z*-score SD<0.2 and BMI percentiles <70% at all time points. This percentile cutoff was arbitrary but made to limit the number of youths who may transition to overweight at a later point. In contrast, WG youth SD was ≥0.2 and had gained ≥38.3 pounds from baseline to the year-two follow-up (1SD above the mean, *M*_weight gain_=25.7±12.6lbs; **Fig1C**). **Fig1D** illustrates that a growth spurt (i.e., height change) was not influencing group dichotomization. Additionally, the criterion captured youth across a range of BMIs (**Fig1E**) and weight classes (**Fig1F** and **Fig1G**). Out of all youth with eligible data, 748 met criteria for the WS_HW_ or WG group (see **Supplemental Materials FigS1** for a flow chart).

**Figure 1.**
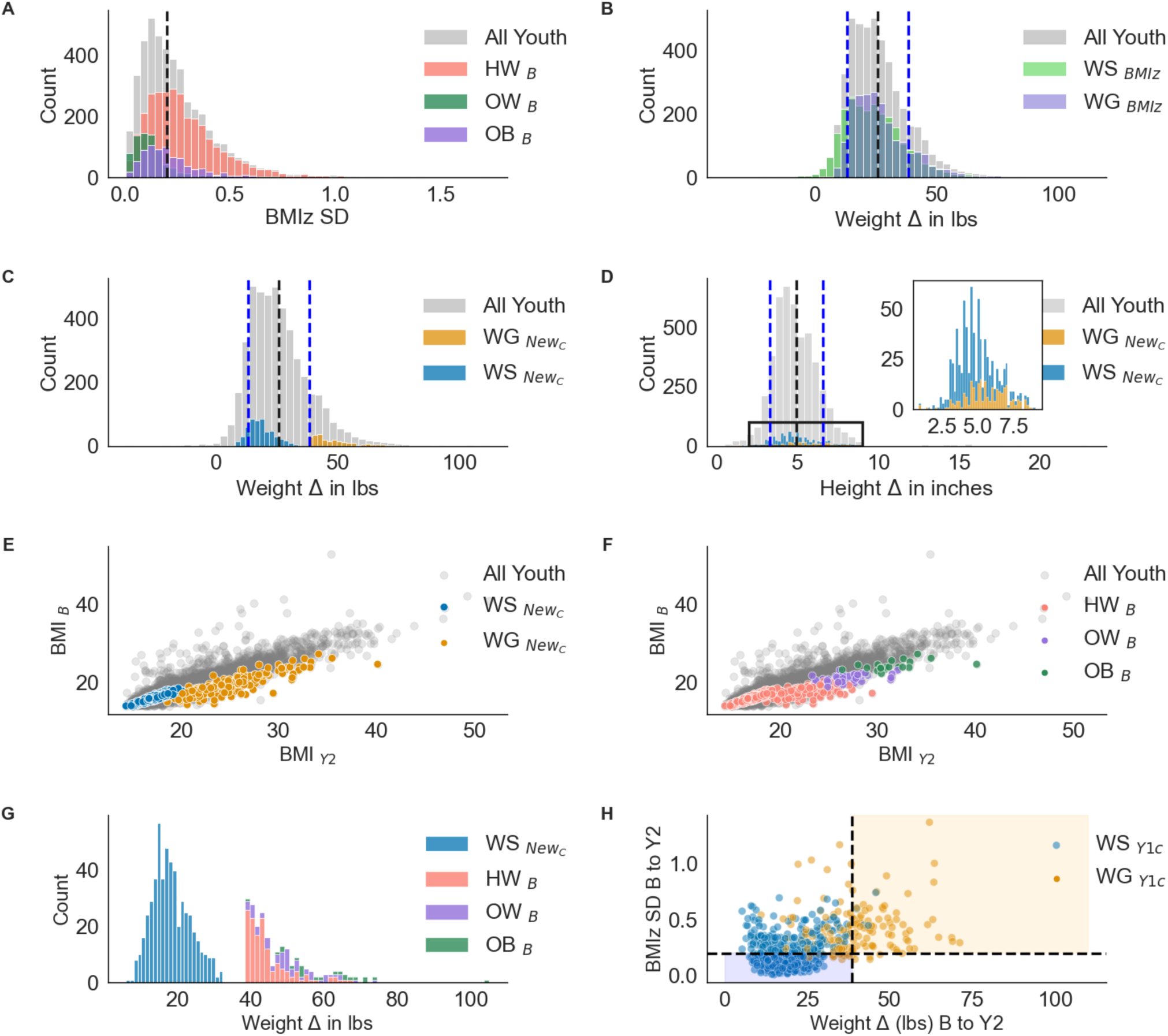
**A)** Distribution of BMI *z-*score (BMIz) standard deviation (SD) for all youth and color coded by baseline(B) weight class (e.g., healthy weight [HW], overweight [OW], obese [OB]). Black dashed line=the clinical cut-off for weight stability as described in the literature.^31, 32^ For Panels **B, C,** and **D**, black dashed line=entire sample mean; blue dashed lines =±1 SD for the entire sample**. B)** The clinical weight stability cut-off does not adequately classify rapid weight gainers (color coded). There were a substantial number of youths who gained more than 1 SD (i.e., >38 lbs) above the mean (25.0 ±12.7 lbs) that met both the weight-stable and weight-gain criterion. **C)** Weight stability redefined (New_C_=WG: >38lbs + BMIz SD >0.2; WS_HW_: BMI*z* SD < 0.2 + BMI %ile <70); Weight-gain averages by group: *M* WS_HW_*=*18.2±5.1 lbs; *M*_WG_=47.6±8.8 lbs **D)** Height-change distributions for the entire sample (*M*=4.91±1.7 inches) and by stability group with the new criteria (WS_HW_ *M* =4.7±1.3 inches; *M*_WS_=5.6±2.4 inches). Height did not confound the weight stability classification. **E, F)** The distribution of raw BMI (unadjusted for age, sex, height and weight) from baseline to year 2 (y2) color coded by weight stability and weight class group. The weight stability criterion selected a subset of the youth across a range of BMIs. **G)** The weight stability criterion color coded by baseline weight status. At baseline, 62% of the WG group were classified at baseline, while 17% remained HW at year 2 but were still classified as weight gainers. **H)** Not all youth identified as WS_HW/_WG in our previous report^19^ met the criteria at year 2 (WG=32.5%, WS_HW_=39%). Y1c=Year 1 classification. Colored boxes=the youth who met Y1 and Y2 criteria, vs. white areas=youth classified previously as WS_HW_ / WG at Y1^19^ but not at Y2. Black dashed lines indicate the cut-off for weight stability defined in this manuscript. Note: The abscissa for all subplots is on different scales.

**Figure 2.**
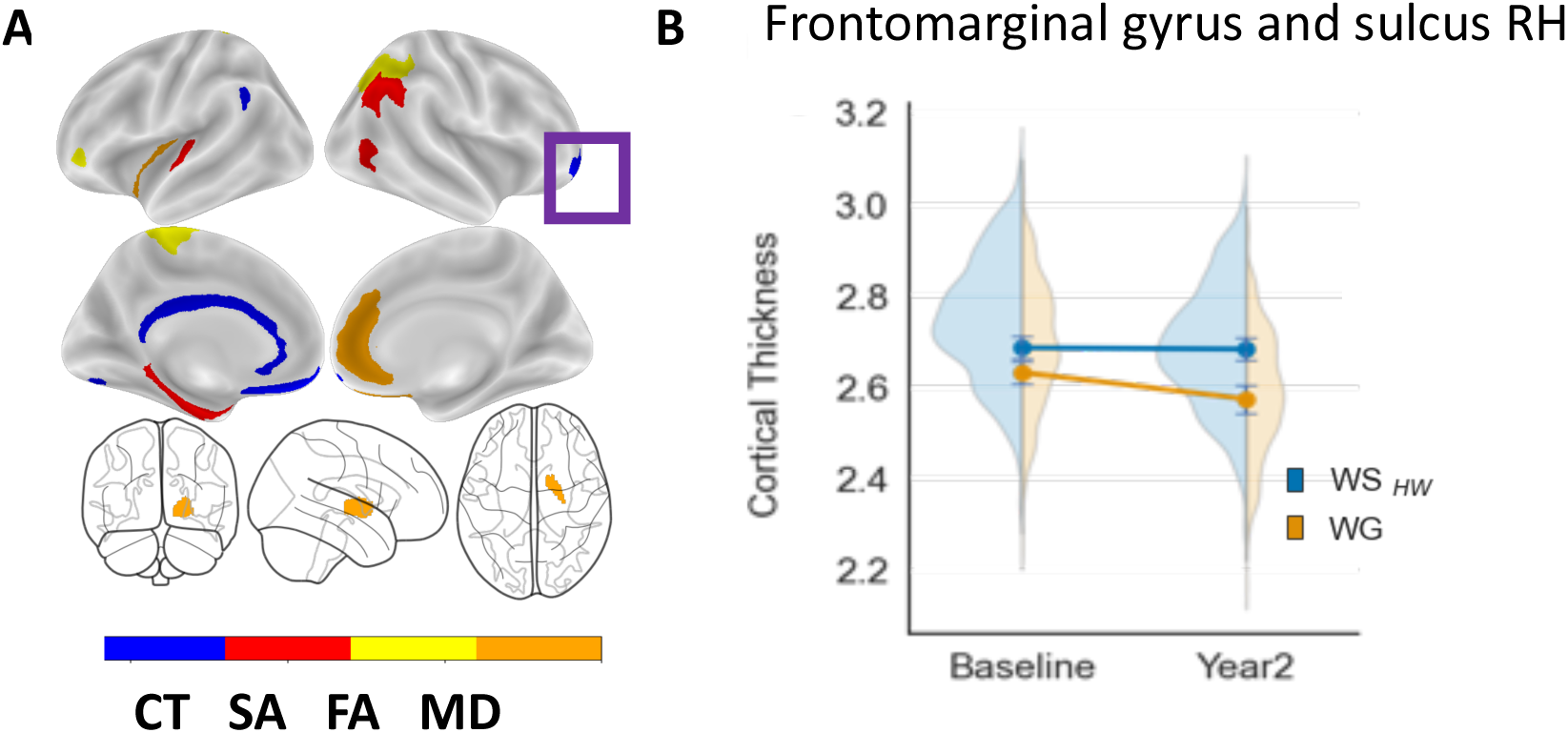
**A)** Visualization of the brain regions by modality associated with weight gain (i.e., >20 lbs) at the one-year period (previously identified in Adise et al., 2021).^19^ Purple box=significant longitudinal change in the frontomarginal gyrus **B)** Frontromarginal gyrus and sulcus change over time. Significant interaction (*p*=0.002) from the mixed model assessing how these regions changed over time weight gain onset. Mixed model effects were independent of age, sex, puberty, race/ethnicity, highest household education, BMI, time, and caregiver report of prenatal exposure to alcohol and tobacco.

**Figure 3.**
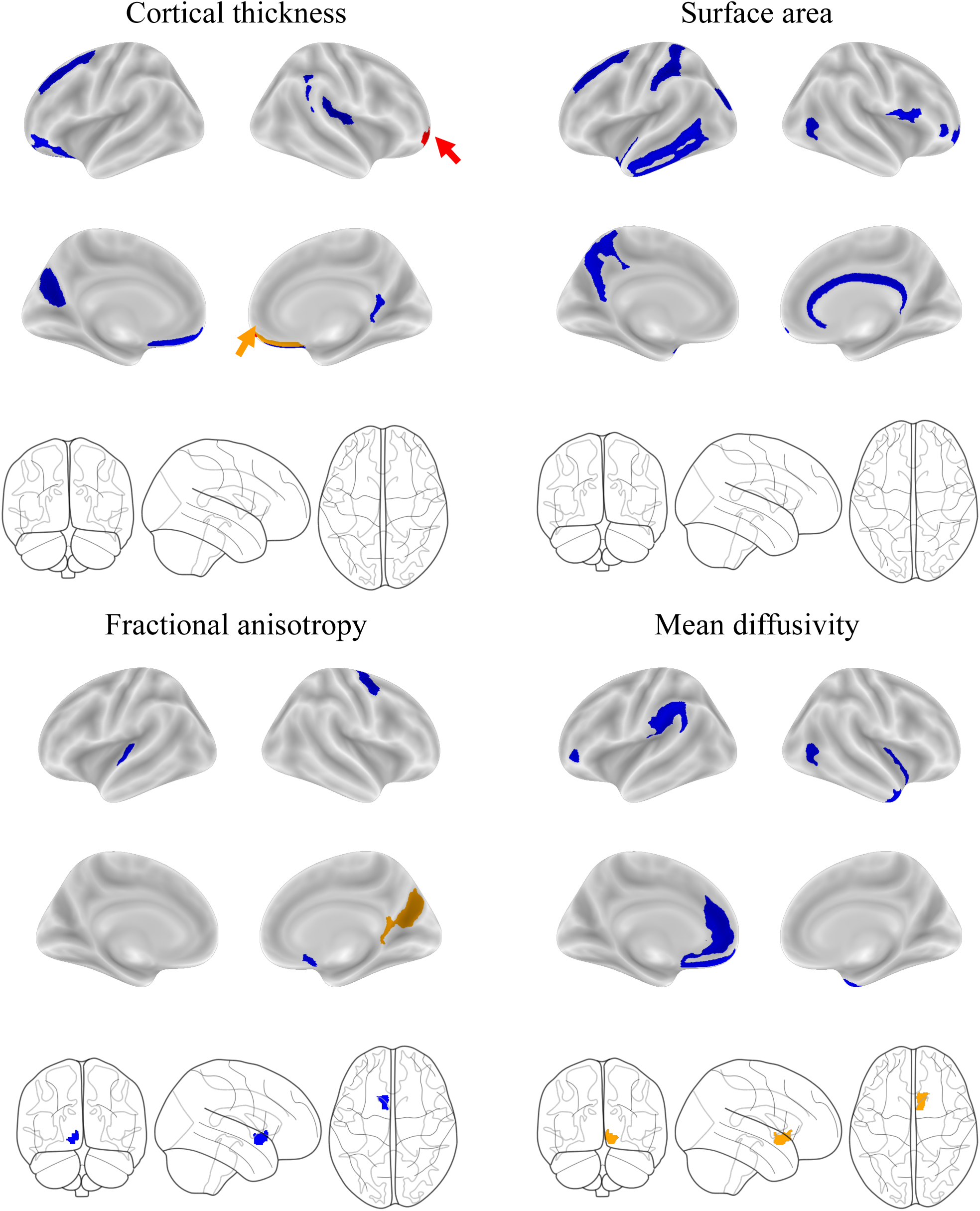
Visualization of the baseline brain features across each modality identified from the elastic-net regression that predicted youth in the WG group after two-years of sustained, rapid weight gain. Color schematics represent significant main effects and interactions (corrected for multiple comparisons for Group and Group by Time) from the mixed model assessing how these regions changed over time weight gain onset. Mixed model effects were independent of age, sex, puberty, race/ethnicity, highest household education, BMI, time, and caregiver report of prenatal exposure to alcohol and tobacco. Blue=no significant change. Orange=Significant main effect of Group. Red=Significant interaction between Group and Time. The arrows highlight areas difficult to see.

Only 32.5% of WG youth and 39% WS_HW_ at the one-year follow-up published previously^19^ were included in analyses here, either because data were not yet available for the two-year follow-up, some youth with one-year rapid weight gain^19^ did not show continued weight gain trajectory over the two-years (**Fig1H, Supplemental Materials TableS2**), or because some youth initially identified as WS_HW_ at the one-year follow-up were no longer WS_HW_ at the two-year follow-up (**Fig1H**).

### Pubertal assessment

Puberty was assessed via caregiver and self-report sex-specific questionnaires. Scores were converted into sex-specific Tanner staging categories^38^ and averaged across caregiver and youth reports (1=Prepubertal, 2=Early puberty; 3=Mid puberty; 4=Late puberty; 5=Postpubertal).

### Demographic assessments

Caregiver-reported child’s race/ethnicity, date of birth, and sex at birth were obtained at the baseline visit (see **Supplemental Materials** for details).

### Prenatal assessments

Caregivers reported preterm delivery (yes/no/refuse), weeks born premature, birth weight, prenatal tobacco exposure before and after pregnancy confirmation (yes/no/refuse), and prenatal alcohol exposure before and after pregnancy confirmation (yes/no/refuse).

### Kiddie schedule for affective disorders and schizophrenia for school-age youth (KSADS)

The KSADS assessed psychiatric illnesses (including eating disorders, such as binge eating, anorexia, and bulimia) via caregiver report. In sum, 32 Diagnostic and Statistical Manual of Mental Disorders-5 (DSM-5) child psychiatric diagnoses were created including codes for present, remission, and lifetime diagnoses and converted into 0 (absence of diagnosis) or 1 (definitive diagnosis). KSAD diagnoses for eating disorders were used as exclusion criteria (see **Supplemental Materials TableS3** for details).

### Cognitive Assessments

Youth completed a modified and original Behavioral Inhibition System/Behavioral Approach System (BIS/BAS) questionnaire, which is used to assess trait-based reward and inhibitory control. Youth also completed the Urgency, Premeditation, Perseverance, Sensation Seeking, and Positive Urgency (UPPS-P) Impulsive Behavior Scale, which assess impulsivity (see **Supplemental Materials Methods and TableS3** for details).

### Neuroimaging acquisition and preprocessing

MRI data were collected with 29 scanners: Details on data acquisition and analyses are published elsewhere^26, 39^. The current manuscript focuses on structural MRI data, collected with the T_1_-weighted, and Diffusion Tensor Imaging (DTI) acquisitions. Cortical data were parcellated with Freesurfer using the Destrieux Atlas (148 regions of interest [ROI]). Volumetric data was parcellated before surface projection using an atlas of 16 ROIs. Structural data consisted of cortical thickness (mean thickness per ROI), surface area (total surface area per ROI), and subcortical volume. Fractional anisotropy (FA) and mean diffusivity (MD) white-matter ROI estimates sub-adjacent to each cortical were extracted from full-shell DTI images. Subcortical estimates for each FA and MD ROI reflect a mixture of both white- and gray-matter estimates.

## Statistics

### Mixed Models ROI Analysis

Multiple linear mixed models were conducted in Python with the pymer4 package^41^ to determine the relationship between weight gain and change in brain structure, while accounting for inter-individual differences. The mixed models corrected for BMI, sex, age, highest household education, race/ethnicity, scan year (i.e., baseline, two-year follow-up), and caregiver report of prenatal exposure to tobacco and alcohol. Models that included FA and MD included a motion estimate, and brain models also included intracranial volume as a covariate. Random effects were modeled to account for variability in scanners across the ABCD Study^®^ sites and within-subject variation. Sibling relations were not included because 98% (n=731) of youth in our analyses were singletons. Dependent variables consisted of ROIs associated with predicting (1) one-year weight gain (n=18 regions)^19^ or (2) two-year weight gain (identified from the elastic-net regressions below) or (3) cognitive assessments.

Categorical variables were dummy coded, with the reference variable set to the largest *n* per category. Pairwise comparisons were corrected for multiple comparisons with the Tukey’s approach. Main effects (WS_HW_ vs. WG) and the group-by-time interactions were corrected for multiple comparisons using the Benjamini-Hochberg approach across each modality (e.g., cortical thickness, surface area, FA, MD, and subcortical volume). Correction was conducted separately for the ROIs associated with one-year and two-year weight gain. Statistical reporting for categorical predictors in the main text, tables, and figures are reported using the *F*-statistic using pymer4’s *anova* function.

### Identification of brain regions associated with weight gain over a two-year period by utilizing an elastic-net regression

A five-fold cross-validation (80% train, 20% test) logistic elastic-net regression was employed with the Brain Predictability toolbox Python package^42^ (see **Supplemental Materials** for more details) to identify ROIs that were indicative of sustained, two-year weight gain in comparison to our previous published findings,^19^ which may only be predictive of short duration, one-year weight gain. Three elastic-net regressions were run to get the independent effects of brain only, nonbrain only (i.e., covariates like age, sex, puberty, race/ethnicity, highest household education, scanner ID), and a model that combined both brain and nonbrain features. Only models that included brain and nonbrain features are reported in the main manuscript, while the results from the additional models are reported in the **Supplemental Materials TablesS6.**

## RESULTS

### Sample Characteristics

The WS_HW_ group consisted of 527 youth (*M*_WS_=18.2±5.1 lbs). The WG group consisted of 221 youth (*M*_W_=47.7±9.1 lbs; **Fig1C** and **Fig1D, Table1**), and 63.3% were classified as having a healthy weight at baseline (**Table1**). Despite rapid weight gain, 17.6% remained of a healthy weight at the two-year follow-up (**Fig1G**).

**Table 1.** Demographics for the Weight Stable (WS_HW_) and Weight Gain (WG) groups. BMI=Body Mass Index; lbs.=pounds; Λ1=change score; *M*=mean; *SD*=Standard deviation. HS=High school; GED=Generalized Education Diploma; BA= Bachelor’s degree. Weight class was determined by the Center for Disease Control’s sex-age-height-weight-specific growth curves.^29^ *p_group_=*significant group differences between WG and WS_HW_. *p_all_* =significant differences between WG, WS_HW_ and the rest of the sample. *p*-values represent significance testing for *t*-tests and chi-squared testing.

|  |  | WS<sub>HW</sub><br>(n=527) | WG<br>(n=221) | *P*<sub>group</sub> | Other<br>(n=3869) | *P*<sub>all</sub> |
| --- | --- | --- | --- | --- | --- | --- |
| Age [*M* (*SD*)] | *Baseline* | 119.6 (7.3) | 121.8 (7.2) | 0.001 | 119.7 (7.4) | 0.001 |
|  | *Y2* | 143.5 (7.5) | 145.8 (7.6) | 0.001 | 143.5 (7.7) | 0.001 |
| Puberty [*M* (*SD*)] | *Baseline* | 1.8 (0.7) | 2.3 (0.8) | <0.001 | 2.0 (0.8) | <0.001 |
|  | *Y2* | 2.5 (0.9) | 3.2 (0.9) | <0.001 | 2.7 (1.0) | <0.001 |
| BMI [*M* (*SD*)] | *Baseline* | 16.3 (0.9) | 19.3 (2.5) | <0.001 | 19.6 (4.0) | <0.001 |
|  | *Y2* | 17.4 (1.1) | 24.7 (3.5) | <0.001 | 21.3 (4.5) | <0.001 |
| Weight Δ in lbs [*M* (*SD*)] |  | 18.2 (5.1) | 47.7 (9.1) | <0.001 | 25.5 (12.1) | <0.001 |
| Height Δ in inches [*M* (*SD*)] |  | 4.7 (1.3) | 5.6 (1.5) | <0.001 | 4.9 (1.7) | <0.001 |
| Sex [*n* (%)] | *Male* | 281 (53.3) | 97 (43.9) | 0.023 | 2058 (53.2) | 0.026 |
|  | *Female* | 246 (46.7) | 124 (56.1) |  | 1811 (46.8) |  |
| Race [*n* (%)] | *White* | 397 (75.3) | 124 (56.1) | <0.001 | 2188 (56.6) | <0.001 |
|  | *Black* | 29 (5.5) | 36 (16.3) |  | 456 (11.8) |  |
|  | *Hispanic* | 52 (9.9) | 41 (18.6) |  | 777 (20.1) |  |
|  | *Asian* | 8 (1.5) | 1 (0.5) |  | 76 (2.0) |  |
|  | *Other* | 41 (7.8) | 19 (8.6) |  | 372 (9.6) |  |
| Highest Household Edu [*n* (%)] | *<HS* | 9 (1.7) | 11 (5.0) | <0.001 | 135 (3.5) | <0.001 |
|  | *HS/GED* | 13 (2.5) | 17 (7.7) |  | 281 (7.3) |  |
|  | *Some College* | 98 (18.6) | 89 (40.3) |  | 976 (25.2) |  |
|  | *BA degree* | 145 (27.5) | 45 (20.4) |  | 1072 (27.7) |  |
|  | *Postgraduate degree* | 262 (49.7) | 59 (26.7) |  | 1405 (36.3) |  |
| Baseline Weight Class [*n* (%)] | *Healthy Weight* | 527 (100.0) | 140 (63.3) | <0.001 | 2388 (61.7) | <0.001 |
|  | *Overweight* |  | 59 (26.7) |  | 716 (18.5) |  |
|  | *Obese* |  | 22 (10.0) |  | 765 (19.8) |  |
| Y2 Weight Class [*n* (%)] | *Healthy Weight* | 527 (100.0) | 39 (17.6) | <0.001 | 2345 (60.6) | <0.001 |
|  | *Overweight* |  | 89 (40.3) |  | 761 (19.7) |  |
|  | *Obese* |  | 93 (42.1) |  | 763 (19.7) |  |

The WG group differed significantly from those in the WS_HW_ group on all demographic variables (age, sex, BMI, puberty, race/ethnicity, parental highest education). Youth in the WG group were two months older (*p*<0.001), had more advanced puberty (*p*<0.001), and consisted of more females (*p*=0.026). Additionally, the WG group had higher percentages of Black and Hispanic youth (*p*<0.001) and lower percentages of parents with advanced education (*p*<0.001; **Table1**).

Mothers of youth in the WG group reported higher rates of short-term prenatal tobacco exposure (i.e., exposure during the first trimester, but discontinued post pregnancy recognition; *n*=40, 18.1%) compared to youth in the WS_HW_ group (*n*=52, 10%; *p*<0.001). Although significantly different, only 7% of youth in the WG group were exposed to prenatal tobacco continuously (i.e., exposure during all three trimesters; *n*=15) compared to youth in the WS_HW_ group (*n*=42, 4%; *p*=0.03). However, youth in the WS_HW_ group were more likely to be exposed to continuous alcohol exposure (*n*=16, 3%), than those in the WG group (*n*=5, 2%; *p*=0.40, **Supplemental Materials TableS4**). Because youth in the WG group differed on these demographic factors, they were controlled for in the subsequent analyses.

*Do regions identified as predictive of one-year rapid weight gain*^19^ *show continued structural change over a two-year period of sustained weight gain?*

Mixed models revealed that while controlling for age, sex, BMI, puberty, race/ethnicity, education, intercranial volume, and caregiver report of prenatal exposure to alcohol and tobacco, there was a main effect of Group (*F*(1, 1092.61)=19.2, *p*<0.001) and a Group × Time interaction (*F*(1, 1241.5)=10.0, *p*<0.001) in the right frontomarginal gyrus and sulcus (**TableS5**). Youth in the WG group had thinner cortices (*M*=2.602±0.025, 95%CI [2.545,2.59], **Fig4A**) in this region compared to those in the WS_HW_ group (*M*=2.684±0.023, 95%CI [2.634,2.736]] and showed greater acceleration of thinning in this region by year two (*M*=2.573±0.03, 95%CI [2.51,2.644]; **Fig4C**) than WS_HW_ youth (*M*=2.682±0.03, 95%CI [2.652,2.274]). For a complete list of main effects and interactions, please refer to **TableS5**.

*Are there underlying differences in brain structure at baseline that can predict group membership (e.g., WS_HW_/WG) two years later?*

A logistic elastic-net regression identified 39 brain ROIs, along with baseline age, baseline puberty, highest household education, and motion, as features that predicted WG group membership two years later (AUC_train_=0.75, Matthews Correlation Coefficient [MCC]=0.38; AUC_test_=0.68, MCC=0.22). In the test set, 17 out of 33 (52%) WG and 81 out of 109 (74%) WS_HW_ youth were correctly identified, while the overall balanced accuracy was 62.5%. The brain features included 10 cortical-thickness ROIs, 13 surface-area ROIs, 5 cortical-FA ROIs, 9 MD ROIs, 1 subcortical-FA ROI, and 1 subcortical-MD ROI (see **FigS2, Table2**). ROIs identified for rapid weight gain over two years were largely non-overlapping (92%, *n*=36) with those identified for one-year rapid weight gain^19^ (**Table2)**. Results for the brain only and nonbrain only (i.e., covariates) are presented in the **Supplemental Materials TableS6.**

**Table 2.** Results for the logistic elastic-net regression showing which baseline features predicted group membership (i.e., WS_HW_ / WG) based on two years of sustained, rapid weight gain. In other words, these sets of features are associated with weight prior to weight onset and may be potential biomarkers of continued weight gain beyond a one-year period. Region of interest (ROI) labels are in accordance with the Destrieux atlas labels. G=gyrus; S=sulcus; L=left; R=right;

| Feature | Beta weight |
| --- | --- |
| *Cortical thickness* |  |
| Frontomarginal gyrus RH <sup>b</sup> | -0.09422 |
| Medial orbito-olfactory sulcus RH | -0.04753 |
| Orbital gyrus LH | -0.03948 |
| Parieto-occipital sulcus LH | -0.06094 |
| Posterior ramus of the lateral sulcus RH | -0.0668 |
| Posterior ventral cingulate gyrus RH | 0.012326 |
| Rectal gyrus LH <sup>b</sup> | -0.01181 |
| Rectal gyrus RH | -0.01684 |
| Sulcus intermedius primus (of Jensen) RH | 0.008321 |
| Superior frontal sulcus LH | -0.02883 |
| *Surface Area* |  |
| Anterior occipital sulcus RH <sup>b</sup> | -0.01295 |
| Frontomarginal gyrus RH | -0.00979 |
| Inferior circular insula sulcus RH | 0.003891 |
| Inferior frontal opercular gyrus RH | 0.038942 |
| Inferior temporal gyrus LH | -0.02406 |
| Lateral orbital sulcus RH | 0.014423 |
| Middle temporal gyrus LH | -0.06602 |
| Pericallosal sulcus RH | 0.083268 |
| Planum polare of the superior temporal gyrus LH | -0.11852 |
| Postcentral sulcus LH | 0.03637 |
| Precuneus gyrus LH | -0.0761 |
| Superior frontal sulcus LH | 0.048763 |
| Superior occipital sulcus and transverse occipital sulcus LH | 0.011407 |
| *Fractional anisotropy* |  |
| Accumbens area LH | 0.029662 |
| Anterior transverse temporal gyrus LH | -0.0033 |
| Parieto-occipital sulcus RH | -0.04135 |
| Posterior ventral cingulate gyrus RH | -0.01394 |
| Subcallosal gyrus RH | 0.044645 |
| Superior part of the precentral sulcus RH | -0.00169 |
| *Mean diffusivity* |  |
| Accumbens area RH | -0.02754 |
| Anterior cingulate gyrus and sulcus LH | -0.07266 |
| Anterior occipital sulcus RH | -0.05326 |
| Long insular gyrus and central sulcus of the insula RH | -0.06025 |
| Rectal gyrus LH | -0.01869 |
| Subcallosal gyrus LH | -0.00035 |
| Superior circular insula sulcus LH | -0.01971 |
| Supramarginal gyrus LH | -0.01258 |
| Temporal pole RH | -0.01419 |
| *Nonbrain features* |  |
| DTI Motion | 0.119316 |
| Some college | 0.093156 |
| Postgraduate education | -0.05622 |
| Baseline Age | 0.11139 |
| Baseline Puberty | 0.249004 |

**Table 2.** Results for the logistic elastic-net regression showing which baseline features predicted group membership (i.e., W<sub>SHW</sub> / WG) based on two years of sustained, rapid weight gain.
| Feature | Beta weight |
| --- | --- |
| *Cortical thickness* |  |
| Frontomarginal gyrus RH <sup>b</sup> | -0.09422 |
| Medial orbito-olfactory sulcus RH | -0.04753 |
| Orbital gyrus LH | -0.03948 |
| Parieto-occipital sulcus LH | -0.06094 |
| Posterior ramus of the lateral sulcus RH | -0.0668 |
| Posterior ventral cingulate gyrus RH | 0.012326 |
| Rectal gyrus LH <sup>b</sup> | -0.01181 |
| Rectal gyrus RH | -0.01684 |
| Sulcus intermedius primus (of Jensen) RH | 0.008321 |
| Superior frontal sulcus LH | -0.02883 |
| *Surface Area* |  |
| Anterior occipital sulcus RH <sup>b</sup> | -0.01295 |
| Frontomarginal gyrus RH | -0.00979 |
| Inferior circular insula sulcus RH | 0.003891 |
| Inferior frontal opercular gyrus RH | 0.038942 |
| Inferior temporal gyrus LH | -0.02406 |
| Lateral orbital sulcus RH | 0.014423 |
| Middle temporal gyrus LH | -0.06602 |
| Pericallosal sulcus RH | 0.083268 |
| Planum polare of the superior temporal gyrus LH | -0.11852 |
| Postcentral sulcus LH | 0.03637 |
| Precuneus gyrus LH | -0.0761 |
| Superior frontal sulcus LH | 0.048763 |
| Superior occipital sulcus and transverse occipital sulcus LH | 0.011407 |
| *Fractional anisotropy* |  |
| Accumbens area LH | 0.029662 |
| Anterior transverse temporal gyrus LH | -0.0033 |
| Parieto-occipital sulcus RH | -0.04135 |
| Posterior ventral cingulate gyrus RH | -0.01394 |
| Subcallosal gyrus RH | 0.044645 |
| Superior part of the precentral sulcus RH | -0.00169 |
| *Mean diffusivity* |  |
| Accumbens area RH | -0.02754 |
| Anterior cingulate gyrus and sulcus LH | -0.07266 |
| Anterior occipital sulcus RH | -0.05326 |
| Long insular gyrus and central sulcus of the insula RH | -0.06025 |
| Rectal gyrus LH | -0.01869 |
| Subcallosal gyrus LH | -0.00035 |
| Superior circular insula sulcus LH | -0.01971 |
| Supramarginal gyrus LH | -0.01258 |
| Temporal pole RH | -0.01419 |
| *Nonbrain features* |  |
| DTI Motion | 0.119316 |
| Some college | 0.093156 |
| Postgraduate education | -0.05622 |
| Baseline Age | 0.11139 |
| Baseline Puberty | 0.249004 |

At baseline, WG youth had thinner cortices (80% of the predictive CT ROIs), decreased MD (100% of the predictive MD ROIs), and decreased FA (67% of the predictive FA ROIs) but greater surface area (64% of the predictive SA ROIs; see Supplemental Materials FigS2 for a visual representation). Baseline age, puberty, and motion during DTI were positive predictors of WG youth two-years later, while household highest education was a negative predictor of group membership.

*Do regions found at baseline to be predictive of* WS_HW_*/WG group membership two-years later show additional longitudinal structural changes over time (i.e., after weight gain onset)?*

Out of the 39 baseline brain features associated with two-year sustained weight gain, only six brain features (all right hemisphere) showed significant group differences in change over the two years (*p’*s<0.001). Mean differences, confidence intervals, and significance are reported in TableS7. The Supplemental Materials contain visual representations of these effects (FigS3-5). When compared to WS_HW_ youth, over the two-years, WG youth had greater reductions in cortical thickness in the right frontomarginal gyrus (aforementioned, *t*(4.4), *p<*0.001, FigS3A) and rectal gyrus (*t*(3.04),*p*=0.001, FigS3D), greater reductions in FA in the right parieto-occipital sulcus (*t*(3.66),*p*<0.001) and posterior ventral cingulate gyrus (*t*(3.31),*p*=0.001), and greater reductions in MD in the nucleus accumbens (t(4.42),p<0.001). Effects were independent of age, sex, baseline puberty, BMI, race/ethnicity, highest household education, intracranial volume, caregiver report of prenatal exposure to alcohol and tobacco, and motion (for FA and MD). No other main effects or interactions were observed (**TableS7** lists effects and interactions, corrected and uncorrected; **TableS8** shows posthoc comparisons). Youth who were overweight/obese at baseline were removed from the analyses to confirm that the prediction model was not being driving by brain/BMI associations. Even with a reduced sample size (*n*_WG_=140), the elastic net identified WS_HW_ versus WG_HW_ youth (AUC_test_=0.76, MCC=0.34, balanced accuracy=0.71, confusion matrix percent correct=67%).

### Behavioral differences between groups

Lastly, to effectively contextualize the behavioral relevance of neurostructural differences between WS_HW_ and WG groups, we conducted a series of analyses to determine how the WS_HW_ and WG groups differed with respect to impulsive behaviors.

*BIS/BAS*: No effects survived multiple-comparisons correction (**TableS9**).

*UPPS-P:* WG youth scored significantly higher on the lack of perseverance subscale (i.e., the tendency to quit when a task gets hard or boring) (*F*(1, 1020.12)=7.6*, p*=0.005, *M*=7.3±0.3, 95%CI [7.11,8.35]) than those in the WS_HW_ group (*M*=7.11±0.3, 95%CI [6.55,7.67]; **TableS10**, **FigS6A**). For positive urgency (i.e., the tendency to respond impulsively to positive affective states), there was a Group × Time interaction (*F*(1, 1195.06)=11.0, *p*=0.002, **FigSC**). At baseline, WG youth scored higher (*M*=9.02±0.3, 95%CI [8.26,9.77]) compared to WS_HW_ youth (M=8.46±0.3, 95% CI [7.71,9.21], *p*=0.03), but no differences were observed at the two-year follow-up (*M*_WSHW_=7.95±0.371, 95%CI [7.18,871], *p*=0.2; *M_WG_*=7.43±0.4, 95%CI [8.4,9.06]).

All main effects and interactions were independent of age, sex, BMI, baseline puberty, race/ethnicity, highest parental education, and caregiver report of prenatal exposure to alcohol or tobacco. No other main effects or interactions were observed (**TableS10**).

## Discussion

Much research has focused on understanding the metabolic consequences of rapid weight gain during infancy, but little is known about this weight gain phenomena later in development. Here, for the first time, we show that (1) there are different trajectories of rapid weight gain (i.e., short, one-year; sustained, two-year), (2) structural variation at 9-to-10-years-old can predict who will experience rapid weight over longer periods of time (two years), and (3) the pattern of brain structural differences between WS_HW_ and WG groups are largely non-overlapping for short-versus long-term weight gain. Further, we show that trajectories of maturational change in some brain regions are altered by two-years of rapid weight gain *after* its onset. These findings are of considerable significance, as they show that individual differences in brain structure may predispose such individuals to sustained unhealthy weight gain over longer periods of time and that brain structure may change as a function of the excessive weight gain itself. Overall, our results attest to the power of studies like the ABCD Study^®^ given its longitudinal design, permitting investigation of competing hypotheses/predictions, and its large demographically diverse sample, allowing for the analytical power to investigate relatively rare phenotypes/outcomes (e.g., rapid weight gain independent of diagnostic criteria for eating disorders).

Adolescence is a developmental period of risk for weight gain due to the effects of puberty on growth^46^ and the emergence of eating disorders.^11, 12^ Our research highlights the importance of studying rapid weight gain during adolescence as we observed phenotypic differences in weight gain (i.e., short vs. sustained) that corresponded with different patterns of structural variation predicting which youth would experience rapid weight gain over a short (i.e., one-year) and longer (i.e., two-year) duration. When compared to one-year weight gain,^19^ two-year weight gain was predicted by more widespread structural variation at 9-to-10-years-old in regions involved with executive functioning (e.g., frontomarginal and rectal gyri, and superior frontal sulcus), reward and appetitive control processing (e.g., nucleus accumbens, anterior cingulate), emotion regulation (e.g., anterior and postrior cingulate gyrus), and working memory (e.g., parieto-occipital sulcus).^45–47^ Decreased available resources (e.g., reduced cortical thickness, decreased FA) across more regions associated with decision-making and food intake, may make it increasingly hard for some youth to inhibit food intake, and thereby result in longer durations of rapid weight gain. Animal studies add support to this theory as neuroinflammation affects the brain prior to weight gain onset^48^ and is modulated by dietary intake.^49^ Therefore, it may be that greater structural variation *prior to* two-year weight gain may be indicative of more neuroinflammation that coincides with increase caloric intake of high fat foods. This corresponds with our observed phenotypic differences suggesting that food intake patterns may differ amongst those with short and sustained weight gain. Additional longitudinal data will continue to add to our knowledge of the neural differences between these weight gain phenotypes. However, combined with our previous work,^19^ this demonstrates the predictive power of brain structure as a potential biomarker for identifying youth who are at risk for gaining weight over a short and relatively longer period of time.

After two-years of weight gain onset, we observed continued structural change in some of the regions identified to be predictive of prospective two-year (but not one-year) weight gain. By 11-to-12-years-old, youth with two-years of rapid weight gain continued to present with significant differences from their WS_HW_ youth counterparts in regions implicated in food intake and obesity^45–47^ such as the frontomarginal and rectal gyrus, parieto-occipital sulcus, ventral gyrus, and nucleus accumbens. Structural variation in these regions may send the wrong signals to the hypothalamus to trigger food intake^49^ that creates a predisposition to and contributes to weight gain maintenance via a cyclical pattern garnered in food intake facilitation and subsequent future weight gain. Because our data only covered a two-year time span, future studies are needed to assess the role of structural variation and maintained rapid weight gain.

Interestingly, no other regions that were predictive of two-year weight gain showed sustained changes after weight gain, nor did baseline regions that were predictive of one-year weight gain. There are several explanations for this: It may be that (1) a two-year period during development is not long enough to observe detrimental consequences of weight gain on brain structure, (2) that differences in brain structure occurred earlier (or will occur later) in development, or (3) regions predictive of one-year weight gain may be indicative of another phenotype of weight gain (i.e., short term) that may be subject to little variation over time. Although the neuroinflammatory effects of weight gain have been established in humans and animals,^49^ its temporal course is not clearly defined. Neuroinflammatory effects on brain structure may be moderated by diet and exercise,^50, 51^ and normative developmental brain maturation and reorganization^52, 53^ may adjust deleterious effects of weight gain on brain structure if the detrimental effects of neuroinflammation can be reversed. While these findings serve as a reference point, additional research is needed to understand how the brain changes over a prolonged period in response to short and sustained (i.e., multi-year) weight gain during this period of maturation.^52^

The literature suggests that one reason for overeating may be rooted in deficits in impulsivity^20, 54^ (i.e., the ability to control urges to eat). Our data supports this possibility as WG youth presented with thinner cortices in regions that are involved with impulsivity (e.g., frontomarginal gyrus and sulcus). Additionally, WG youth were more likely to act impulsively during a positive mood (i.e., positive urgency) and scored higher on the lack of perseverance subscale of the UPPS-S. Together, this suggests that rapid weight gain may be partly explained by deficits in impulsivity regarding food intake decisions. However, the ABCD Study^®^ did not measure objective food intake or how decisions may change in the presence of food. Therefore, more research is needed to understand how structural changes in inhibitory control regions relate to weight gain and impulsive food choices over time.

### Strengths and limitations

To our knowledge, this was the first study to assess how the brain changes *after* two-years of rapid weight gain in regions showing structural variation *prior to* weight gain. Because youth did not meet diagnostic criteria for binge eating or other eating disorders, these findings have relevance for understanding the brain’s role in rapid weight gain development, maintenance, and its deleterious neurological and cognitive effects. However, the ABCD Study^®^ did not collect measurements of emotional overeating or objectively measured food intake, which limits the inferences that can be made from the brain and weight-gain associations. While the WG youth differed significantly from those in the WS_HW_ on several key demographics, these variables were added as covariates in the mixed models, though, we lacked statistical power to further explore these potential associations. Moreover, due to sample-size limitations, we were not able to investigate longitudinal phenotypic and neurological differences between youth who had short versus sustained weight gain. Fortunately, future releases of the ABCD Study^®^data may afford larger samples of these two phenotypes so that we can further understand how youth who experience shorter durations of weight gain (and then plateau) differ from those with sustained weight gain. While the elastic net chose the best model that was predictive of weight gain, it is possible that there are other regions that are also important but that were not selected for inclusion in the model. Lastly, information about parental weight, a strong predictor child obesity risk, was not collected by the ABCD Study^®^.

## Conclusion

Adolescence is a period in development at risk for excess weight gain. Despite growing obesity trends, little is known about the causes and effects of rapid weight gain and its temporal nature. The current study sheds light on the predictive power of the brain as a biomarker for identifying youth with rapid weight gain trajectories who may present with subclinical eating disorders but do not yet meet diagnostic criteria. Moreover, a large percentage of youth were of a healthy weight *prior to weight gain*, but still exhibited patterns of structural variation that were associated with weight gain prediction and maintenance. These findings add to the literature suggesting that overeating may produce neuroinflammatory effects that cause brain structure variation *prior to weight gain*, suggesting that changes in the brain may be an early marker of obesity later in life. Two-year rapid weight gain was preceded by structural variation in brain regions associated with inhibitory control, emotion regulation, and appetitive control, and these differences were maintained over a two-year period. This suggests that structural variation in these regions may be important not only for initiation but for continuation of rapid weight gain trajectories throughout adolescence. Further, these findings add to the growing body of literature aimed at understanding the causes and consequences of rapid weight gain during adolescence.

Follow-up studies are needed to examine how these brain structures continue to differentially relate to rapid weight gain trajectories in adolescence.

## Acknowledgements

SA wrote the manuscript and performed the analyses with the guidance of ES, AM, MH. SA conceptualized and curated the data with the help of ES. ES and MH provided funding for the project as part of the ABCD Study consortium. EK, AM, and SZ contributed to interpretation and data analysis. All authors contributed feedback, and read, and approved the final manuscript. The ABCD Study consortium investigators designed and implemented the study and/or provided data but did not participate in the analyses or writing of this manuscript.

